# An end-to-end model of active electrosensation

**DOI:** 10.1101/2024.10.22.619741

**Authors:** Denis Turcu, Abigail Zadina, L.F. Abbott, Nathaniel B. Sawtell

## Abstract

Weakly electric fish localize and identify objects by sensing distortions in a self-generated electric field. Fish can determine the resistance and capacitance of an object, for example, even though the field distortions being sensed are small and highly-dependent on object distance and size. Here we construct a model of the responses of the fish’s electroreceptors on the basis of experimental data, and we develop a model of the electric fields generated by the fish and the distortions due to objects of different resistances and capacitances. This provides us with an accurate and efficient method for generating large artificial data sets simulating fish interacting with a wide variety of objects. Using these sets, we train an artificial neural network (ANN), representing brain areas downstream of electroreceptors, to extract the 3D location, size, and electrical properties of objects. The model performs best if the ANN operates in two stages: first estimating object distance and size and then using this information to extract electrical properties. This suggests a specific form of modularity in the electrosensory system that can be tested experimentally and highlights the potential of end-to-end modeling for studies of sensory processing.

## 2 Introduction

Weakly electric fish sense their environment by emitting electrical fields known as electric organ discharges [1]. The electric field around the fish associated with an electric organ discharge, which we refer to as the EOD, consists of a basal EOD, which is the field that would exist in empty water, plus the electric field induced due to nearby objects, called the electric image, which appears as a distortion in the basal EOD. Nearby objects with electrical resistances higher than the surrounding water (e.g. rocks) result in less EOD-induced current flow near the object, producing a local decrease in the amplitude of the EOD. Living objects (e.g. small invertebrates that are prey for the fish), on the other hand, have lower electrical resistances than water and hence increase field amplitude. Living objects also have sizable electrical capacitances, which alters the temporal waveform of the EOD.

The outcome of EOD signal processing is the remarkable ability of weakly electric fish to spatially localize objects and characterize their properties (including size, shape, and electrical resistance and capacitance) in the dark, based solely on information extracted from their EODs [2, 3, 4]. While the importance of localizing objects and determining their size and shape is obvious, the unique ability of electric fish to discriminate electrical properties is likely to be of special importance for foraging by aiding the fish in finding preferred prey [5]. The object-induced perturbations of the EOD that support electrosensation are typically small and are highly sensitive to distance (decreasing as 1/distance^4^) and to object size (increasing as radius^3^). This limits the distances over which the fish can determine object properties to the multi-cm range. The species studied here, *Gnathonemus petersii*, emits pulsatile EODs of ∼ 1 V amplitude and ∼ 300 *µ*s duration. Behavioral studies suggest that microvolt changes in EOD amplitude and sub-microsecond temporal distortions of the EOD waveform can be detected by the fish [2, 3, 4, 5]. Although the initial stages of electrosensory processing have been intensively studied [6, 7, 8], how information contained in subtle perturbations of the EOD is transformed into behaviorally meaningful representations of object location and identity remains largely unknown.

The EOD is sensed by approximately 1, 000 electroreceptor organs distributed across the fish’s body surface, each of which contains two classes of receptors known as A- and B-cells [9]. A- and B-cells encode different features of the EOD (see Section 3) and project to separate regions of the electrosensory lobe (ELL), the first stage of electrosensory processing in the fish’s brain [10, 11, 12, 13]. Projections from these two regions, the medial zone (MZ) for A-cells and the dorsolateral zone (DLZ) for B-cells, converge in the midbrain and are subsequently processed within an interconnected network of brain regions including the optic tectum, thalamus, telencephalon, and cerebellum [14, 15, 16].

To investigate the processing of EOD signals, we begin by constructing models of electroreceptor responses and electric field generation that allow us to simulate the elctrosensation of objects with varying locations, sizes, and electrical properties. We then use these large simulated datasets to train a variety of ANN architectures to simultaneously localize objects and identify their electrical properties, a task solved by the fish during foraging.

## 3 Results

### 3.1 Measuring and modeling electroreceptor responses

Our first goal was to develop a model of the sensory information transmitted by A- and B-type electroreceptors. Prior electrophysiological recordings have shown that A-type receptors are primarily sensitive to changes in EOD amplitude, whereas B-type receptors respond to both amplitude and waveform changes, but a precise description of the stimulus features encoded by A- and B-type receptors is lacking [10, 12, 13, 17]. To address this, we recorded responses in both the MZ and DLZ to a large set of simulated EODs designed to mimic objects with different resistances and capacitances (Fig 1 A). In these experiments, fish were paralyzed, blocking the action of the electric organ, so both the basal EOD and the distortions in it were generated artificially, triggered by electrophysiological measurement of EOD command signals. Delivered fields were recorded to verify that they matched the desired waveforms (Sup Fig 8). These stimuli generated prominent field potentials which we recorded with microelectrodes positioned at matched somatotopic locations in the MZ and DLZ. Based on previous results, we used the amplitude of the first negative peak of the LFP (henceforth called the LFP amplitude) as a proxy for the activity of individual electroreceptor afferent nerve fibers [18, 19, 20]. Distorting the basal EOD evoked large and reliable changes in LFP amplitude in both zones (Fig 1 B). We report EOD distortions and sensory responses as percentage differences from the basal EOD or the response to it.

**Figure 1:**
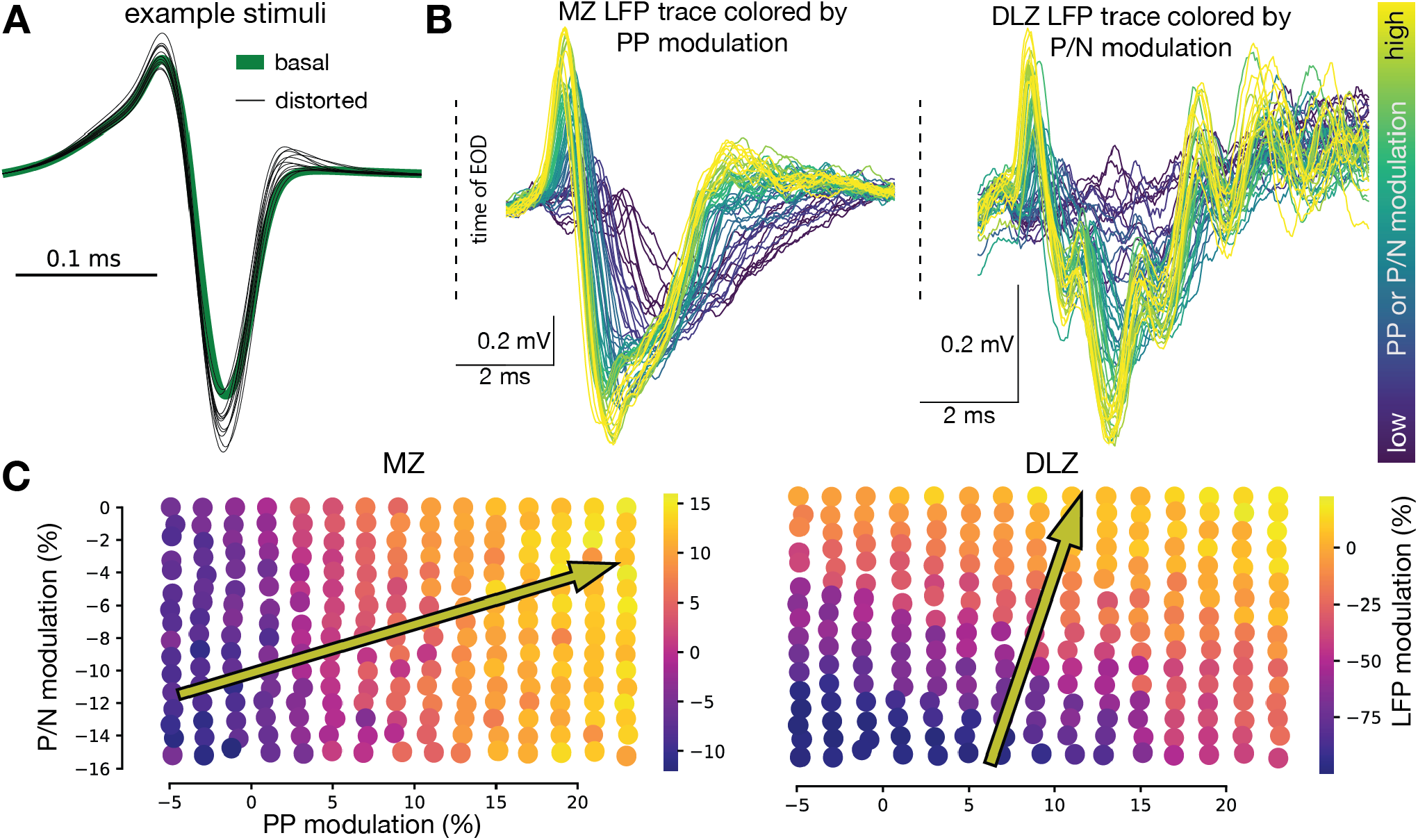
Responses to resistive and capacitive stimuli in the medial and dorsolateral zones of the ELL. **A** Examples of delivered EODs. The basal EOD (green) is plotted behind individual examples (gray) that include distorted EODs with different amplitudes and waveform shapes, simulating the effects of objects with different electrical properties. **B** Example LFP responses to delivered stimuli from a single fish. The color for each trace reflects the PP amplitude modulation for MZ (left) and the P/N ratio modulation for DLZ (right). Dashed black line marks the timing of the EOD. Traces for each of the 56 different distorted stimuli delivered in this example experiment are shown. **C** Summary of LFP responses for all stimuli in the PP and P/N feature space, color-coded by the MZ response (left) and the DLZ response (right). Data from a single fish from an experiment in which 240 different distorted stimuli were delivered. Arrows indicate the directions of the gradients of MZ and DLZ responses in the feature space.

Previous studies [5, 21] have characterized distortions of the EOD due to resistive and capacitive objects in terms of changes in the peak-to-peak amplitude (PP = P+N) and the positive-to-negative peak ratio (P/N) of the EOD waveform (Fig 1 A). Plotting LFP amplitude as a function of the PP and P/N values of the corresponding stimuli, we found that responses in the MZ depend primarily on the PP value of the stimulus (Fig 1 C, left), while those in the DLZ depend more on P/N (Fig 1 C, right). However, the gradients of the measured responses are not truly aligned (see Section 6 for alignment details) with either of these two features (arrows in Fig 1 C).

To provide a better description of the measured LFP responses, we constructed a model based on convolving the distorted EOD waveforms with two filters, one for the MZ and another for the DLZ (Fig 2 A). Because we characterize electrosensory responses by the amplitude at a single time point (the magnitude of the negative peak in the LFP), the convolution took the form of a projection, i.e. a product of the stimulus waveform and the filter, integrated over time. We also added an offset parameter to this sum. We determined the filter shapes and offsets by minimizing the squared difference between the prediction of the filter model and the data across EOD stimuli. Artificial stimulus noise was included as a regularizer in this minimization, with the appropriate noise value chosen by cross-validation (Sup Fig 9 A). The resulting model explains 92% of the variance across single trial responses (Fig 2 C) and provides interpretable convolutional filters that are robust across experiments (Fig 2 B). The filter shapes suggest that A-type receptors weigh and sum the three peaks of the EOD waveform (Fig 2 B, left), while the B-type receptors are sensitive to temporal features of the EOD waveform, including the slope and timing of the zero-crossing (Fig 2 B, right).

**Figure 2:**
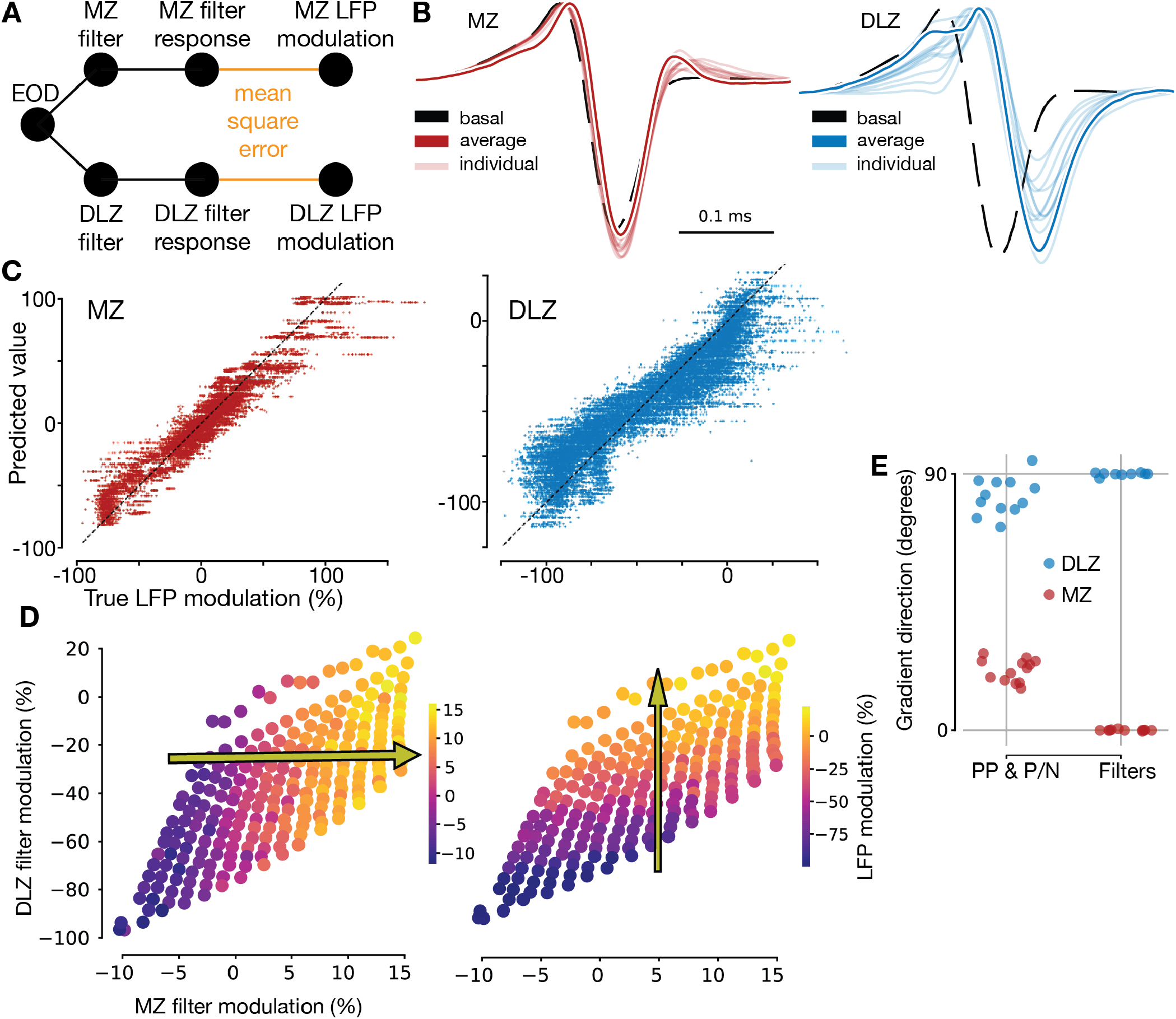
Convolutional filter model of MZ and DLZ responses. **A** Schematic of the model. **B** Filters obtained from fitting the data for both filter types. Base EOD (black), average filter (solid color), and individual experiments filters (light color) are shown. **C** Performance of the filter model in predicting the single-trial LFP response. **D** Summary of LFP responses for all stimuli in the filters feature space, color-coded by the MZ response (left) and the DLZ response (right). Data from a single fish, same experiment shown in Fig 1 C. Arrows indicate the direction of gradients of the responses in the feature space. **E** Summary across experiments of the alignment of the LFP responses with the PP & P/N features (left) and the filters features (right) (n = 13, MZ; n = 12, DLZ).

As in Fig 1 C, we plotted the experimental responses as a function of stimulus features, only now using the projections of the stimuli onto the two filters as our axes (Fig 2 D). The MZ and DLZ responses are better aligned (Fig 2 E) with the features extracted from our model (arrows in Fig 2 D) than with the PP and P/N feature space (arrows in Fig 1 C). We also compared our filters with the results of a principal component analysis (PCA) on the set of experimentally delivered stimuli. Two principal components (PCs) explain most of the variance across stimuli (Sup Fig 9 B), matching the number of sensory cell types. Moreover, the first two PCs resemble the filters extracted by our model (Sup Fig 9 C, compare to Fig 2 B), with the main difference being that the PCs are required to be orthogonal by construction.

### 3.2 From objects to EODs

To study the neural computations underlying realistic electrosensory tasks, we need to expand beyond the experimental data to compute responses from the entire electroreceptor array for objects that vary in their electrical properties, size, and location. Detailed numerical models have been developed to compute the spatial patterns of object-induced modulations of the basal EOD [22, 23]. While these models have proven extremely useful for studies of electrosensory systems [22, 24, 25, 26, 27, 28], they have two significant drawbacks for our purposes. First, they are static methods, meaning they do not simulate objects with capacitive properties. Second, they are computationally intensive [29, 30], making them poorly suited for generating the large amounts of simulated sensory input required for training ANN models. An alternative is approximate analytic models [31, 32] or electric circuit models [33], but these do not provide the full flexibility needed for our purposes. We therefore developed a field model framework (see Appendix A) that can quickly and flexibly generate realistic EOD patterns. Our framework captures the spatial geometry of the fish and objects (Fig 3 A), using the field model fitted to data in [34]. It captures the EOD distortions due to both resistive and capacitive properties of objects (Fig 3 B) by solving the dipole distortion problem [31, 32] in a computationally efficient way tailored to this system (Appendix A). It can also reproduce the spatial pattern of an object’s electrical image on the body of the fish (Fig 3 C). This framework can simulate many fish-object conditions at approximately 50 times real-time on a personal computer.

**Figure 3:**
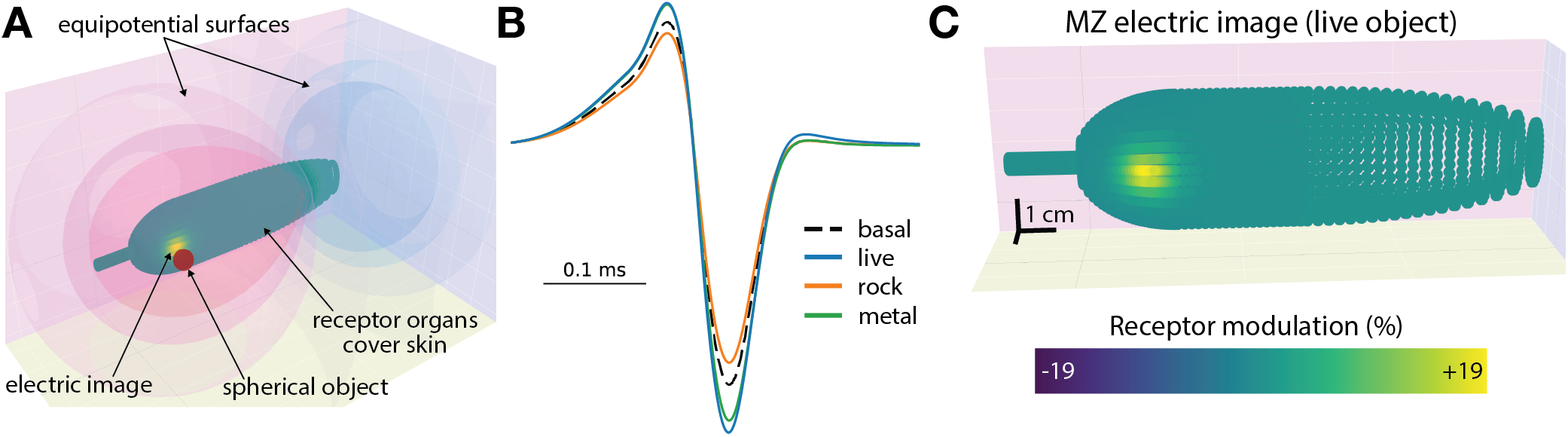
An efficient electric field model for simulating effects of resistive and capacitive objects. **A** 3D visualization of the fish near a spherical object in the aquarium. The fish is covered in model electroreceptor organs and an electric image induced by the object is shown. Equipotential surfaces around the fish during the EOD capture the funneling effect due to the shape of the fish. **B** Example distortions of the EOD due to objects of different electrical properties. The basal EOD is shown in dashed-black. Purely resistive objects distort the amplitude of the EOD, either increasing (distortion due to metal object with small resistance in green) or decreasing (distortion due to rock object with large resistance in orange) the amplitude. Living objects with low resistance and capacitive properties distort both the amplitude and the waveform of the basal EOD (blue). **C** Close-up of the electric image on the skin of the fish visible in A. Modulation is shown as percentage of the basal signal. Individual simulated receptors are visible as green circles on the skin of the simulated fish.

### 3.3 Characterization of object electrical properties

It has been assumed that the electrosensory system derives the resistance and capacitive properties of objects by combining input from A- and B-type receptors, possibly in the midbrain [14, 15]. However, this process has not been studied directly with neural recordings, so it remains unclear how the signals conveyed by A- and B-type receptors support this computation. We therefore examined this process using a modeling approach. We began by training a small, feedforward ANN to extract resistive and capacitive properties of a 2.5 cm spherical object centered at a fixed distance of 2.25 cm from the fish, based on simulated input from a single electroreceptor organ on the skin containing both A- and B-type receptor cells.

We presented the ANN with A- and B-cell inputs to a range of resistances and capacitances that would likely be encountered by a fish (Fig 4 A). We chose the range and the logarithmic spacing of the resistances and capacitances we simulated based on previous experiments [2, 3, 5]. The network successfully extracts these electrical properties with good accuracy, especially for capacitance (Fig 4 B,C). The ANN can also extract resistance and capacitance on held-out individual trials, when provided with the experimental LFP waveforms recorded in the MZ and DLZ as input (prediction error on held-out data was below 5% from true capacitance or resistance values).

**Figure 4:**
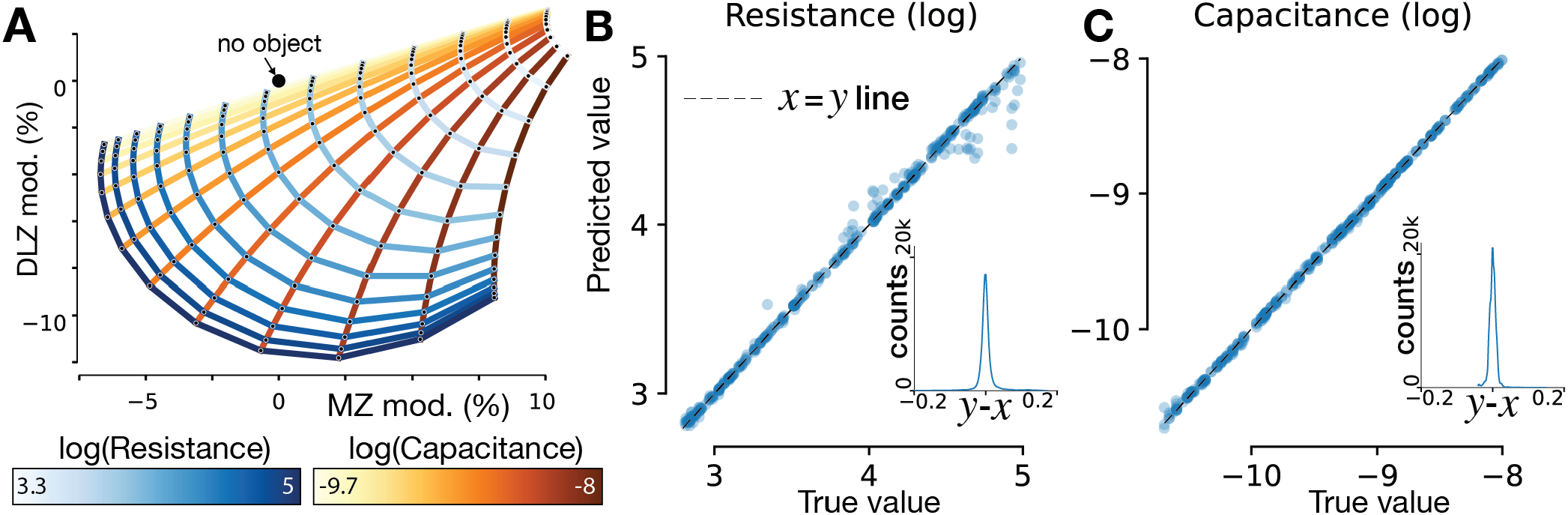
Electric properties for objects of fixed location and size. **A** Objects with different resistances and capacitances occupy different regions of the feature space defined by the MZ and DLZ filters. The size of the spherical object (1 cm) and distance from the fish (0.5 cm) were held fixed. Lines of constant resistance (blue palette) and constant capacitance (orange palette) are shown. The origin in modulation space corresponds to no object present. Resistance and capacitance in all figures are reported as the base 10 log of these quantities in Ω or *F*. **B** Performance of ANN extracting the resistance of different objects with fixed spatial properties. **C** Equivalent to **B** but for capacitance.

### 3.4 Object localization and characterization

The ANN model described in the previous section extracted the capacitance and resistance of an object of fixed size and at a fixed distance. Foraging fish must solve a more complex task, determining the 3D location and size of an object, as well as its electrical properties. The difficulty of this problem can be illustrated by plotting MZ-DLZ feature maps (Fig. 4 A) as a function of the distance to and size of the object generating the EOD distortion (Fig 5 A,B). To obtain these results, we simulated EOD distortions due to objects with varying resistances and capacitances at different distances from the fish (Fig 5 A) and for objects of varying size (Fig 5 B). We chose distance and size values that are typically encountered in experiments [35, 36, 37, 38, 39]. Fish-object distance and object size have a large effect on EOD distortions. From Appendix A, equation 7, it follows that the feature space scaling with inverse distance follows a polynomial of degree 4 for objects within a body length of the fish, and the feature space scaling with object radius follows a polynomial of degree 3. These scalings have a dramatic effect on the performance of the models we now consider.

**Figure 5:**
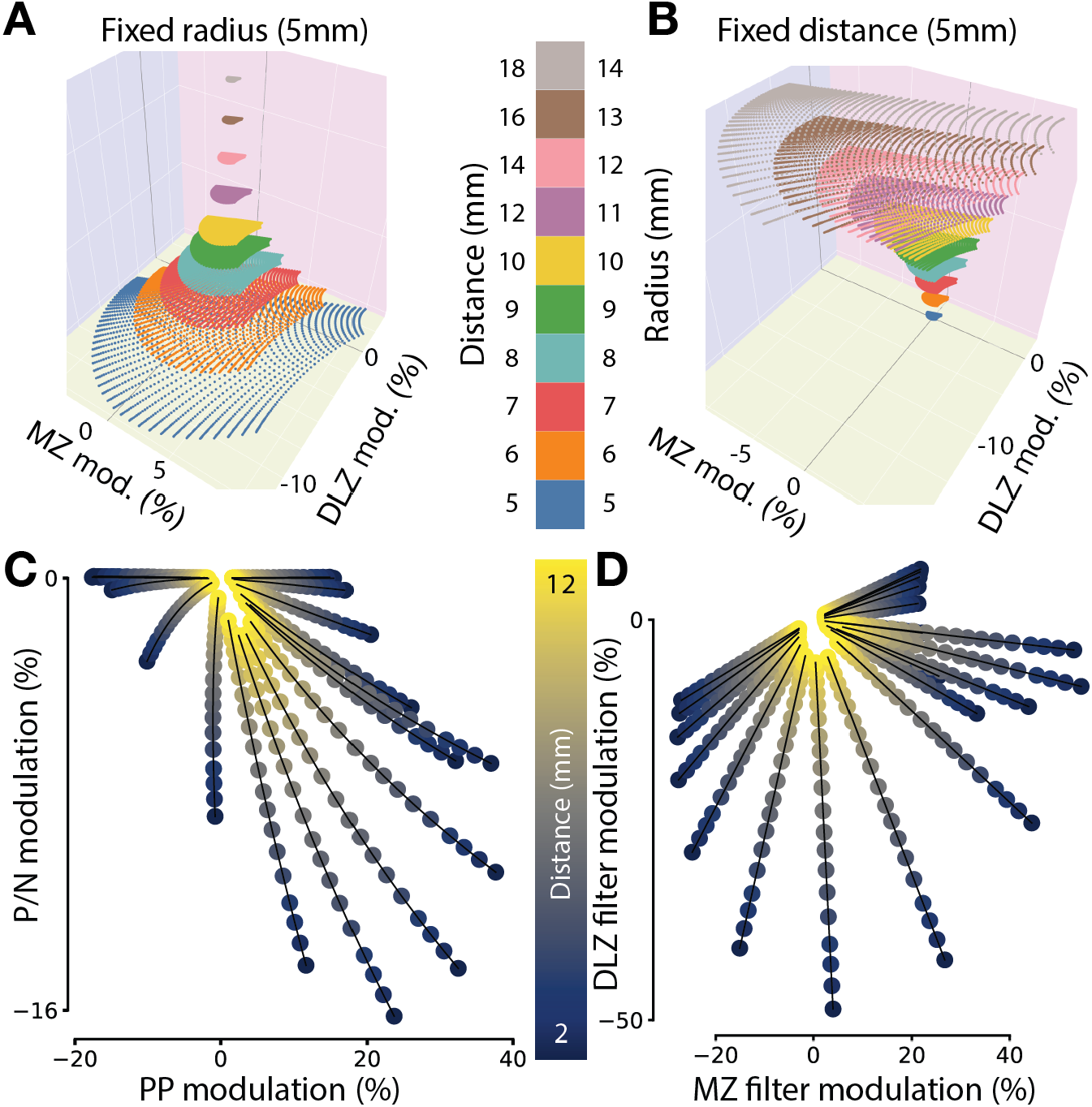
Feature space for active electrosensation. **A** Multiple feature spaces formed by the modulations of the MZ and DLZ filters due to objects with varying resistance and capacitance, with each horizontal plane (different colors) corresponding to a different distance from the fish. Object size and lateral location are fixed. The feature space shrinks by a degree-4 polynomial with inverse distance. **B** Similar to **A**, except each horizontal plane corresponds to the a different object radius, with the distance and lateral location fixed. The feature space increases by a degree-3 polynomial with radius. **C** Amplitude and waveform modulations of the stimulus for objects of fixed size, but of different electrical properties and distances from the fish. Individual points are colored by the distance to the fish and represent distinct combinations of resistance and capacitance. Points corresponding to fixed electric properties but different distances defining electric color lines. In the PP and PN feature space, the electric color lines are not straight — average *R*^2^ for all 20 shown lines is 0.93 with standard deviation 0.16. **D** Similar to **C**, but in the filters feature space. The electric color line is perfectly straight in this space — average *R*^2^ for all 20 shown lines is 1.00 with standard deviation < 10^−11^.

Previous work defined an “electric color line” to capture distance effects [40, 41]. In this work, the EOD modulations produced by an object with fixed electrical properties and fixed size, but located at different distances from the fish, were plotted in the PP and P/N feature space (Fig 1 C). It was noted that points corresponding to specific objects at different distances lay on approximately straight lines. Along the corresponding electric color line an object can be perceived as having the same “color” independent of how far away it is from the fish, in analogy to visual colors with a range of physical characteristics appearing similar. Our electric field model replicates this result, showing that the ratio between the qualitative PP & P/N features fall roughly on a line for the same object when the distance to the fish is varied, but this “line” has some curvature (Fig 5 C). This is because the features PP & P/N are not defined by linear operations on the stimulus. Our electroreceptor model is linear, and the electric color line in the filter (as opposed to PP & P/N) feature space is truly linear (Fig 5 D).

We modeled the extraction of both spatial and electric object properties end-to-end using the physics model to generate electrosensory data and the electroreceptor model to provide the sensory input. Prior work indicated that electrical properties are best encoded in the responses of the most modulated receptors, e.g. those close to the peak response across the skin surface (see electric image on skin in Fig 3 A). On the other hand, spatial properties are encoded in spatial features of the electric image across the fish’s skin, such as its 2D location and overall width and height [42, 43, 38]. Thus, in this case, we modeled receptors across the full surface of the fish (not a single receptor as for the ANN above). To accommodate this array of detectors, we used a more sophisticated variant of an ANN, a convolutional neural network (CNN), to extract object properties. The spatial convolutional filters of the CNN integrate information across the skin array, and the CNN processes this information in sequential stages of convolutional and feedforward layers. We reasoned that CNNs would be suitable for this task on the basis of their success in vision tasks. We tested performance of CNNs with varying numbers of layers and parameters to cover the model space from underparameterized to overparameterized and ensure robustness of results (Section 6).

We first trained the end-to-end CNN models to extract the electrical properties of simulated objects from sensory input, with no specific information about the spatial variables provided during training (no spatial information was included in the loss function, although spatial properties affect the sensory input). The resulting networks perform poorly on extracting electrical properties (Fig 6 A) compared to the previous results when distance to and size were constant (Fig 4 B, C). This is not surprising given the high degree of sensitivity of the feature space to distance and size (Fig 5 A,B). Interestingly, properties of the feature space also account for the structure of the errors in these networks. For resistance, the models distinguish between large value and small values, effectively performing a binary classification. This can be attributed to the fact that scaling of the feature space when distance and size vary preserves angles, but not magnitudes. This allows models to use linear decision boundaries that pass through the origin of the feature space (Fig 4 A) for classification. Based on the lines of equal resistance, splitting the predictions into large and small values results in better than chance performance. For capacitance, the same principle applies, but the equal-capacitance lines are not as favorable for linear decision boundaries passing through the origin, so performance is worse than for resistance.

**Figure 6:**
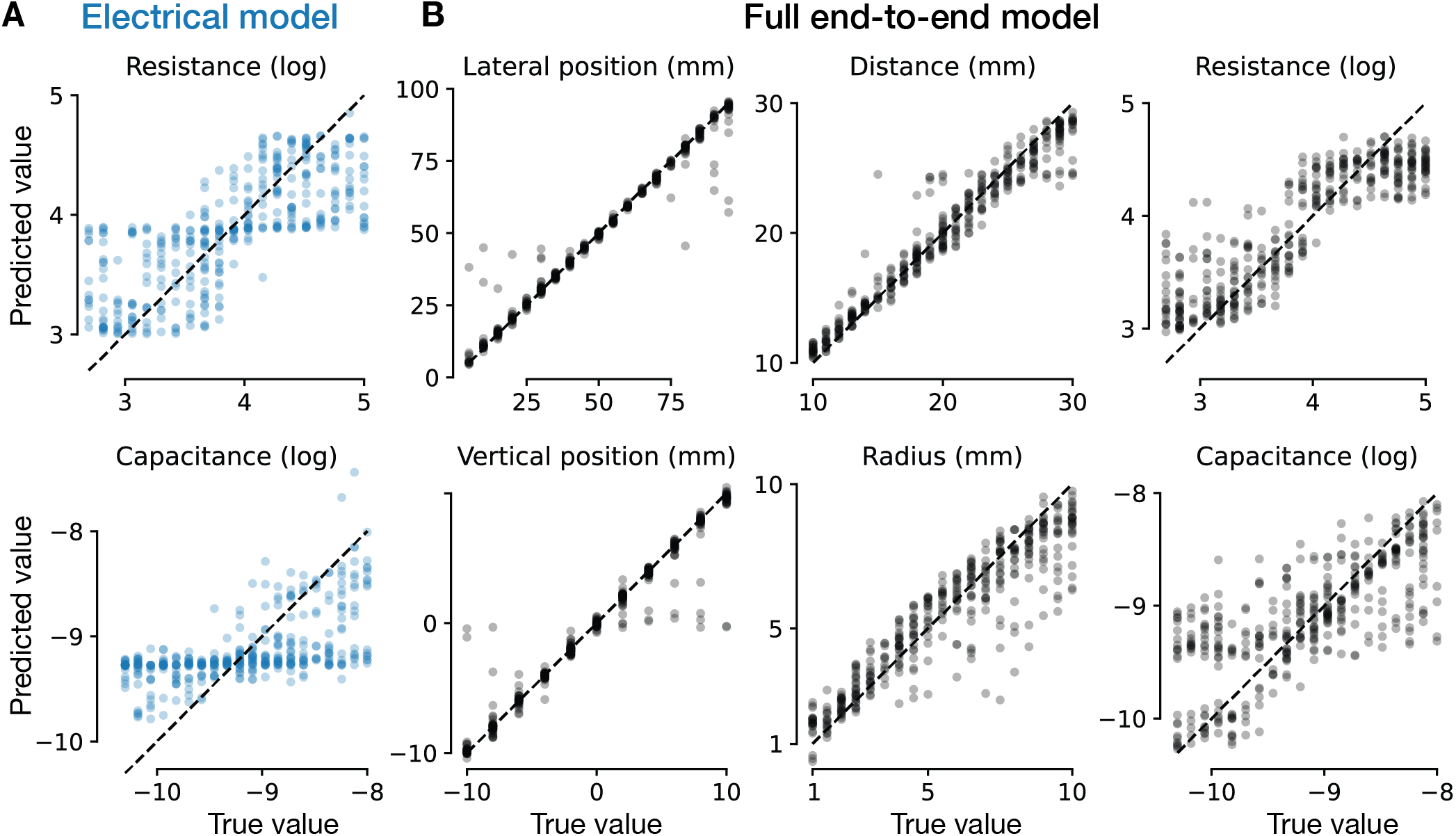
Object localization and characterization by feedforward neural network models. All results reported here are based on cross-validated trials that were not part of training dataset. **A** Performance of spatial-unaware models on extracting the resistance (top) and capacitance (bottom) from the sensory input. This performance shows a binary classification bias of the models extracting the resistance, due to the scaling rule that preserves angles, but not distances in the feature space. The scaling rule angle-preserving effect impairs capacitance performance, explained in part by orientation of the equi-capacitance lines in Fig 4 A). Example performance for one model is shown. The average (± standard deviation) *R*^2^ for each property across *n* = 10 models was: resistance – 0.621±0.008, capacitance – 0.353±0.019. **B** Full end-to-end CNN models’ performance on spatial and electrical properties of simulated objects. Example performance for one model is shown. The average (± standard deviation) *R*^2^ for each property across *n* = 10 models was: lateral position – 0.938 ± 0.017, vertical position – 0.925 ± 0.017, distance – 0.919 ± 0.027, radius – 0.825 ± 0.019, resistance – 0.785 ± 0.021, capacitance – 0.542 ± 0.018.

We next examined if this problem could be solved by training the end-to-end CNN models on the full task, including both spatial and electrical information during training. The resulting models perform well on spatial localization on the extended range of simulated objects chosen to match typical experiments (Fig 6 B left & center). However, they show the same structured errors for the electrical properties (Fig 6 B right) as the electrical-only models, albeit with somewhat improved performance. We reasoned that, even when a CNN is able to extract spatial properties, it may not be able to fully use this information to solve the problems raised by the severe scaling of the feature space, which makes extracting electrical properties hard. In addition, given the steep dependence of the scaling, the CNN’s estimates of spatial properties may not be accurate enough to provide sufficient robustness. In fact, both of these effects contribute to network performance.

To address the problem of using spatial information to inform electrical feature extraction, we combined a CNN trained to determine the spatial features of an object with the ANN that we used previously to extract purely electrical properties (Section 3.3), with one new wrinkle. The small, downstream ANN was pre-trained to learn appropriate multiplicative rules to scale its electrical feature space on the basis of object distance and radius values (Section 6.3). Then the CNN fed its extracted distance and radius values into the downstream ANN, which then applied the learned scaling rule and extracted electrical properties. This hybrid model slightly improved performance for capacitance extraction, but did not improve resistance performance (Fig 7 A). Nevertheless, the downstream ANN implements the scaling rules successfully and, importantly, it accurately extracts the electrical properties of simulated objects with widely varying locations and sizes (Fig 7 B) when the true values of distance and size were fed into it to drive the scaling. This network’s performance is comparable with behavioral performance of fish throughout the same orders of magnitude and for similar logarithmic scaling of resistance and capacitance values that were tested in previous experiments [2, 3]. This indicates that the downstream ANN requires accurate estimates of distance to and size of the object to perform well, more accurate than the CNN model can extract from a single EOD. Fish likely use multiple EODs to localize and characterize objects; they have been observed to emit EODs at rates up to 80 Hz when inspecting objects [44, 36]. This suggests that they might use multiple samples to sharpen their spatial estimates not only to improve spatial localization, but also to better judge the electrical properties of objects.

**Figure 7:**
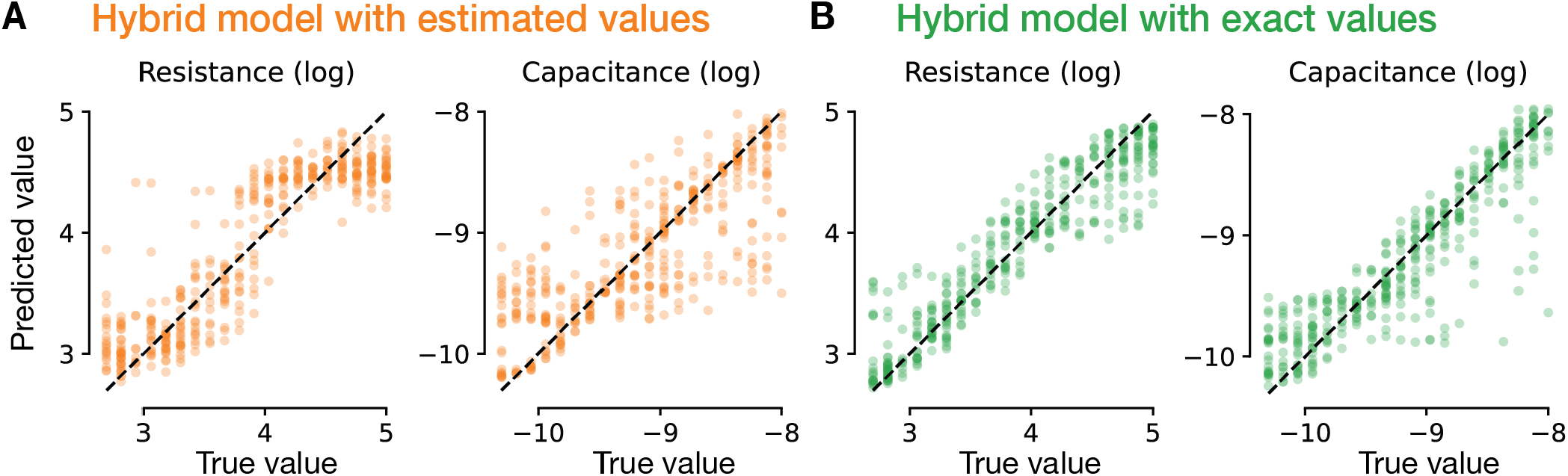
Characterization of object electrical properties by hybrid models. **A** Performance of a hybrid CNN-ANN model with internal spatial estimates from the CNN fed to an ANN that has learned the scaling rule of the feature space and extracts the resistance (left) and capacitance (right) from the sensory input. The average (± standard deviation) *R*^2^ for each property across *n* = 10 models was: resistance – 0.779±0.017, capacitance – 0.597±0.031. **B** Performance of the trained ANN component of the hybrid model when it receives the true spatial values on extracting the resistance (left) and capacitance (right) from the sensory input. The average (± standard deviation) *R*^2^ for each property across *n* = 10 models was: resistance – 0.857 ± 0.019, capacitance – 0.797 ± 0.09. All results here are based on cross-validated trials that were not part of training dataset.

**Figure 8:**
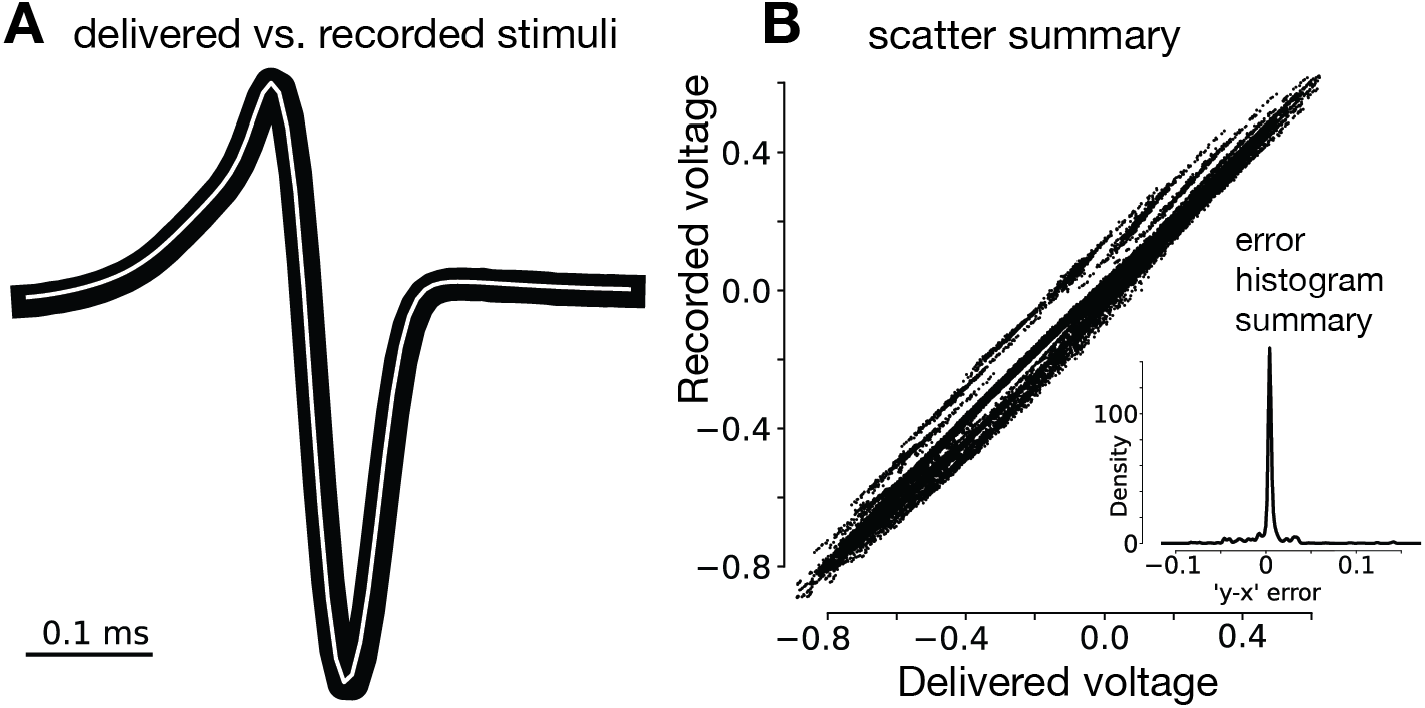
Delivered and recorded stimuli. **A** Example of the delivered stimulus (black) and the recorded stimulus (white) for verifying the accuracy of the simulated EOD waveforms. **B** Summary of scatter points for all stimuli, where each scatter point represents the value of the stimulus at a given time point. Inset marks the histogram of errors of the summary scatter plot.

**Figure 9:**
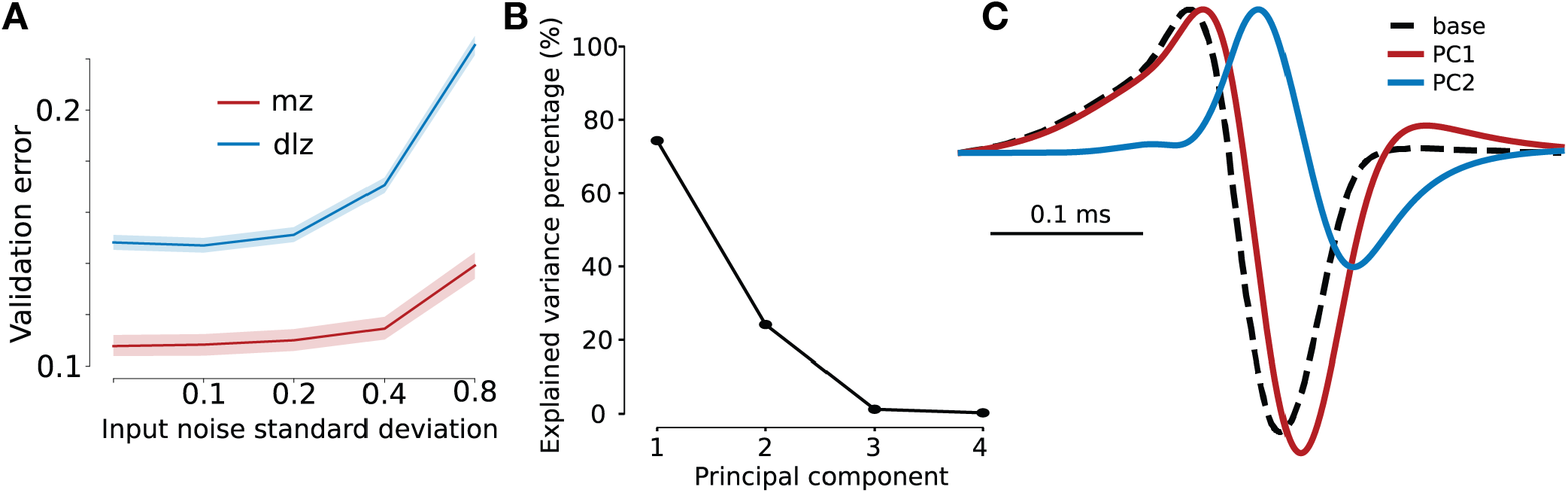
Mormyromast model validation and stimuli PCA. **A** Validation error of the model on held-out single-trial LFP data. This instructs the choice of the noise level in the model. **B** Explained stimulus variance by the first four principal components. **C** The first two principal components of the delivered stimuli.

## 4 Discussion

We have constructed an end-to-end model for extracting spatial and electric properties of objects from EOD signals. In these models, sensory transduction of the EOD was based on stimulus-filters extracted from neural responses evoked by A- and B-type electroreceptor afferents to simulated EODs mimicking the effects of objects with a range of properties. The extracted filters resemble the principal components of the sensory data, correspond to orthogonal directions in a feature space, and generate features that fall along a straight line for objects of fixed resistance and capacitance but varying distances from the fish. These desirable characteristics suggest that the A- and B-type receptors are well adapted to the requirements of electrolocation.

We found that extracting both electric and spatial properties of an object over a range of values is a difficult task for artificial networks, as it must be for the neural circuitry of the fish. The difficulty arises primarily from the high degree of sensitivity of the electrical feature space to distance and size [45]. These spatial aspects scale the electrical feature space, which allows for categorization of objects but makes determining resistance and capacitance values difficult. We solved this problem by pre-training a network to implement distance- and size-dependent scaling operations. Using object distance and radius as an input to scale the feature space to a consistent size simplifies the task that the ANN must solve to extract the resistance and capacitance of the simulated object. Extracting all of the properties of an object, both spatial and electrical, independently is difficult due to strong space-electrical interactions. Pre-training about the nature of these interactions, in particular the existence of a multiplicative scaling of the electrical feature space by distance and radius, resolves this difficulty. It is critical, however, that the internally computed distance and radius be accurate because the scaling is highly sensitive to spatial properties. This insight suggests that an internally driven multiplicative operation could be implemented within downstream brain regions that process electrosensory information. In addition, the idea of using internal computations to drive multiplicative scaling could map onto other systems, such as vision, audition or in artificial intelligence applications, that must deal with scaling effects. For example, internal scaling of images to a standard size on the basis of brain-derived distance estimates could be a useful strategy in vision [46, 47].

Do the brains of electric fish implement anything resembling the two-stage computation described above? Electrophysiological studies of active electrolocation in *Gnathonemus petersii*, as well as other species of African pulse-type electric fish, have primarily focused on the first stage of electrosensory in the ELL [19, 48, 49]. In contrast, very little is known about the neural representation of object size, distance, or electrical properties, which presumably emerge at higher stages of electrosensory processing. Anatomical tracing suggests that information from the MZ and DLZ is fused into a single somatotopic map in a midbrain structure known as the torus semicircularis [14, 15, 16]. Studies of the jamming avoidance response in the South American wave-type electric fish (a behavior that allows fish to avoid electric interference from conspecific EODs), recorded from individual midbrain neurons that combine input from amplitude- and phase-coding pathways (similar to the A- and B-type receptor pathways discussed here) [50]. However, little is known about the potential roles of such neurons in object processing [51]. Midbrain neurons project to “higher” electrosensory processing stages in the optic tectum, cerebellum, and thalamus and also send projections back to the ELL via the preeminential nucleus [14, 52]. This latter pathway has been shown to adaptively shape ELL responses to looming and receding objects in South American weakly electric fish [53]. Our work suggests that signals corresponding to spatial and electrical properties of objects may be processed, at least initially, in separate modules and only later combined. A goal for future studies is to combine the end-to-end modeling approaches developed here with multi-area electrophysiological recordings, ideally in freely swimming fish, to characterize where and how representations of object electrical properties, size, and distance are formed.

## 5 Funding and Acknowledgements

This work was funded by grants from the National Science Foundation (NSF IOS-2115007 to N.B.S.), National Institutes of Health (NIH NS075023 and NS118448 to N.B.S. and L.F.A), the Kavli Foundation (L.F.A. and D.T.), the Gatsby Charitable Foundation (GAT3708 to L.F.A. and D.T.), and the Boehringer Ingelheim Fonds (D.T.). The funders had no role in study design, data collection and analysis, decision to publish, or preparation of the manuscript.

We thank Salomon Z Muller, Federico Pedraja, Avner Wallach and members of the Sawtell and Abbott labs for helpful discussions.

## 6 Methods

### 6.1 Experimental model and subject details

#### 6.1.1 Animals

Male and female wild-caught Mormyrid fish of the species *Gnathonemus petersii* were used in these experiments (fish were 7–12 cm in length, of unknown age, and sex was not specifically selected for). Fish were housed in 60 gallon tanks in groups of 5–20. Water conductivity was maintained between 70–150 *µ*S both in the fish’s home tanks and during experiments. All experiments performed in this study adhere to the American Physiological Society’s Guiding Principles in the Care and Use of Animals and were approved by the Institutional Animal Care and Use Committee of Columbia University.

#### 6.1.2 Method details

##### 6.1.2.1 Surgical procedures

For surgery to expose the brain for recording, fish were anesthetized (MS:222, 1:25,000) and held against a foam pad. Skin on the dorsal surface of the head was removed and a long-lasting local anesthetic (0.75% Bupivacaine) was applied to the wound margins. A plastic rod was cemented to the anterior portion of the skull to secure the head. The posterior portion of the skull overlying the ELL was removed. Gallamine triethiodide (Flaxedil) was given at the end of the surgery (∼ 20 *µ*g/cm of body length) and the anesthetic was removed. Aerated water was passed over the fish’s gills for respiration. Paralysis blocks the effect of electromotoneurons on the electric organ, preventing the EOD, but the motor command signal that would normally elicit an EOD continues to be emitted at an average rate of 2 to 5 Hz.

##### 6.1.2.2 Electrophysiology

The EOD motor command signal was recorded with a Ag-AgCl electrode placed over the electric organ. The command signal is the synchronized volley of electromotoneurons that would normally elicit an EOD in the absence of neuromuscular blockade. The command signal lasts about 3 ms and consists of a small negative wave followed by three larger biphasic waves. Onset of EOD command was defined as the negative peak of the first large biphasic wave in the command signal. Recordings of local field potentials were made with low resistance (< 5 MΩ) glass microelectrodes filled with 2M NaCl. Signals were recorded and filtered at 3–10 kHz (Axoclamp 2B amplifier, Axon Instruments) and digitized at 20–40 kHz (CED power1401 hardware and Spike2 software; Cambridge Electronics Design, Cambridge, UK). For most experiments, recordings were made simultaneously from the MZ and DLZ by placing two electrodes in somatotopically matching locations in the granular layers of the two zones. Somatotopic location and depth within the ELL was judged based on LFP responses to the EOD motor command and to local electrosensory stimuli delivered by a hand-held dipole electrode that could be positioned over various regions of the skin.

##### 6.1.2.3 Electrosensory stimulation

Simulated EODs designed to mimic objects with different resistances and capacitances were delivered using a stimulus isolation unit (A-M systems, model 4100) in constant current mode connected to a pair of carbon rods (2 mm diameter, 4 cm length) placed lengthwise (∼ 1 cm distance from the skin) on either side of the head of the fish. Current amplitude was adjusted for each fish such that the baseline stimulus evoked LFPs of ∼ 70% maximal amplitude. A baseline stimulus consisting of an EOD waveform measured from a discharging fish with no object present was delivered for ∼ 30 minutes before delivering the set of perturbed stimuli. All stimuli were triggered at a brief delay (4.5 ms) following the fish’s spontaneously emitted EOD motor commands.

Three sets of simulated EODs, containing 240, 99 and 56 stimuli, were designed to approximately cover a small rectangular grid in the PP and P/N modulation space. One set of stimuli was used for each fish. Stimuli were delivered in random order in 1–2 bouts containing 10–15 repetitions of a distorted EOD separated by 5–10 repetitions of the baseline stimulus.

### 6.2 Alignment of LFP responses with features

The alignment of mormyromast LFP responses with the feature directions, summarized in Fig 2 E and indicated by arrows in Fig 1 C and Fig 2 D, was computed as the average direction of the slope of the responses in the feature space. For each feature in a feature pair, we computed the slope of the LFP response with respect to that feature’s modulation, while maintaining the other feature approximately constant by binning its modulation values. We used 6 bins for the results reported in Fig 2 E, but results were similar when using other number of bins from 4 to 8. Using more bins resulted in too few samples per bin, and using fewer bins broke the assumption that the other feature was approximately constant. We computed the gradient direction of the alignment as the average of the slopes across the bins.

### 6.3 CNN details

We generated electrosensory data using our field model framework to train the ANN models. The electrosensory dataset contained objects with six properties (3D location, radius, resistance, capacitance) that were independently varied on a grid of values, with between 11 and 32 possible values for each property. The simulated values were selected to cover the distributions of values typically used in experiments for all six properties. The training dataset contained approximately 40 · 10^6^ simulated objects around the fish, and the validation dataset contained approximately 10 · 10^6^ objects. All ANN training was performed in Python using the PyTorch library [54] and the PyTorch Lightning library [55].

The CNN used in Section 3.4 has the same structure as AlexNet [56, 57]. We explored a variety of specific architecture sizes, varying the number of parameters from approximately 200 · 10^3^ to 40 · 10^6^. We varied the number of parameters by changing the number of layers, channels, and neurons. These hyper-parameters ranged from 2–5 convolutional layers with 8–128 channels and 2–4 feedforward layers with 64–5120 neurons. We also applied MaxPool layer to the first and last convolutional layers, and dropout layers with a 0.5 dropout rate to the feedforward layers and trained networks using either *ReLU* or *T anH* activation functions. We used either the Adam and SGD optimizers with learning rates ranging from 0.0001 to 0.02. We used the mean squared error loss function computed on the six objects properties to train the networks. We trained the CNNs to convergence on training data, typically for 50 epochs with a batch size varying from 2, 000– 35, 000. Batch size was chosen to maximize the amount of data that fit in CPU and GPU memory for a training step, according to the network size. We used the validation dataset to monitor the performance of the networks during training, and to select the best model for testing.

The hybrid ANN architecture is only slightly more complex than the CNN. In particular, it uses two separate CNN heads applied to the sensory input — the head receiving the electric image as input and extracting spatial properties of the object (i.e. the same CNN described above), and the head receiving the electric image along with spatial properties, provided either by the spatial head or externally, and extracting electrical properties of the object.

The electric head of the ANN has a much simpler structure than the spatial head, with only one fixed spatial convolutional filter applied to the input, followed by 2–3 feedforward layers with 5–20 units. In between the fixed spatial convolutional and the feedforward layers, this head receives the spatial properties (distance and radius) of the object, and processes them independently to compute a scaling factor for the electrical feature space. The scaling factors are learned by the network, by learning the coefficients of two separate polynomial functions that scale the features with distance and radius. The feature space is scaled by these polynomials before the feedforward layers, and the network is trained to extract the electrical properties of the object from the scaled feature space. The fixed spatial convolutional filter, combined with a MaxPool layer on the whole skin, has the role of finding the most modulated receptor across the skin, and focusing the network on the most informative features for electrical property extraction. This head of the CNN is coupled with the spatial head to extract all object properties.

### 6.4 Code availability

Code is available for each component of this work. The electroreceptors model code is available at github.com/DenisTurcu/efish-receptors-model, the field model framework is available at github.com/DenisTurcu/efish-physics-model, and code for the localization and characterization models is available at github.com/DenisTurcu/efish-characteriation.

## 7 Supplementary figures

### A Electric Field Model

We introduce a model that can generate electrosensory data processed by the fish during active electrolocation. Our model is based on previous work [31, 32, 34], and includes certain extensions that make it suitable for our investigations. With this framework, we can simulate approximately 500 discharges every second, about 10–100 times more than fish typically emit in free behavior. As such, this is suitable for generating fast and accurate electrosensory data for investigating active electrolocation.

#### A.1 Field model details

Electric potential measurements in the environment of weakly electric fish set the basis for modeling the electric field generated by the fish using the EO in their tail. [23, 24] measured the potential on an array of electrodes surrounding the fish and used a numerical method, boundary element method (BEM), to model the electric field. Other studies have used numerical methods such as BEM or finite element method (FEM) to model the electric field generated by weakly electric fish [22, 25, 26, 27, 28] because these methods are accurate and can simulate desireable features of the problem, such as realistic shapes and different conductivity properties for the water, insides of the fish, skin of the fish, and objects in the environment. These numerical methods are broadly designed to solve partial differential equations, such as Poisson’s equation for electrostatics that is of interest for this work, but they suffer from demanding large computational costs across domains of application [58, 59, 60], despite many efforts devoted to speeding up the computations [61, 62, 63]. Additionally, these methods are designed to solve a static problem, but, to fully capture both resistive and capacitive effects of nearby objects, the electric field model should be dynamic.

An analytic field model based on the electric field generated by the fish is more suitable for our investigations. [34] fitted a static multipole model based on data collected by [23, 24] that captures the electric potential surrounding the fish during a discharge. While [34] refers to the electric sources as “charges”, they based their model on the previous work of [31], where the sources are referred to as “currents”. The latter is more appropriate for this system because the fish and its environment are conductive media, where electric charges can move freely instead of remaining stationary. This allows us to account for the effect of the water conductivity on the electric field generated by the fish [31]. Additionally, we can control the temporal waveform of the discharge via the source currents’ amplitude during an EOD to capture the dynamic effects of both resistive and capacitive object properties [31]. These analytic models assume that the fish is electrically transparent with respect to water, but the length of the fish and of the distribution of current sources mimics the field outside of the fish, including the previously coined funneling effect [64]. As such, we use the formalism from [31] in this work, and adapt the model fitted by [34] to suit our investigations.

We model the EOD as *n* + 1 pulse current point sources and sinks, collectively called sources, placed on a segment along the length of the fish, based on the multipole model fit by [34]. The sources are distributed uniformly along the whole length of the fish, *L*, on the mid-line of the fish. As an example, we assume a straight fish lying along the positive 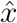− axis, and whose tail is at the origin. The sources are located at 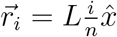 for *i* ∈ {0, …, *n*}. Each source has an associated base magnitude *m*_*i*_, defined as *m*_0_ = −1 and 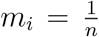 for *i* ∈ {1, …, *n*}. At most times, the sources are inactive, but during the EOD pulse they become active, being multiplied by an appropriately scaled waveform *I*(*t*) = *I*_*o*_*f*(*t*), where *f*(*t*) is the normalized unperturbed EOD. This ensures that at all times, the net current generated by the fish is 0, since current point sources and sinks cancel out. The current flowing through one of the sources is then given by *I*_*i*_(*t*) = *m*_*i*_*I*_*o*_*f*(*t*).

The electric field in an infinite, homogeneous and isotropic conductive medium due to a source, *i*, can be computed using the current density 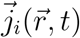at a location 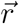 in the medium and the conductivity of the medium, *σ*_*w*_ for water in this case:

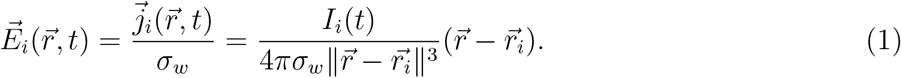

Assuming the fish is electrically transparent with respect to water, the electric field at any point in the medium is given by the superposition of each point current’s electric field:

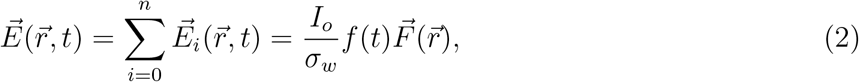

where 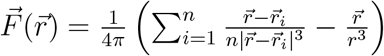.

In this model, fish generate a current flow in their conductive environment, water, that gives rise to an electric field distribution during the discharge. Electric charges must be moving through the medium to generate the current flow. Therefore it is not obvious that we can simply model the discharge waveform by controlling the temporal amplitude of the discharge with a temporal function *f*(*t*), as described above and implied in [31]. Previous multipole-based models of the EOD either have assumed that we can, with brief discussion of this potential problem [31] or have considered a static model analyzing the amplitudes of stimuli only [34]. To motivate the steady-state assumption, we use that a charge distribution inside a conductor decays over a timescale *τ* = *ε*/*σ*. For water with *σ*_water_ ≈ 100*µ*S/cm and *ε*_water_ ≈ 10^−9^F/m, the timescale *τ*_water_ ≈ 10^−7^s is much shorter than the duration of the EOD, approximately 10^−3^s. Therefore, the temporal waveform of the discharge can be controlled by the function *f*(*t*).

#### A.2 Object polarization and dipole distortions

Objects placed in mediums with non-zero electric field distribution, and with different electrical properties than their own, become polarized and distort the original electric field. Here, we discuss the polarization of objects placed in water, close to weakly electric discharging fish, and the field distortions they create. [31] introduced this model for investigating object distortions in the electric field generated by weakly electric fish, with an integral solution to solve for the electric field distortion due to the object. [32] solved the integral problem using Fourier analysis for the wave-type weakly electric fish, *Apteronotus*. Both of these studies have used material electrical properties, i.e. conductivity and relative permittivity of the object and water, to simulate object distortions, but experiments with weakly electric fish often use artificial objects with known macroscopic electrical properties, i.e. resistance and capacitance.

Here, we adapt these previous models to our investigations of the pulse-type weakly electric fish, *G. petersii*. First, we found that the complete Fourier analysis solution from [32] is not applicable to the pulse-type EOD of *G. petersii*, for many realistic choices of object electrical properties, because the harmonic series does not readily converge in these scenarios. Therefore, we combine the Fourier analysis solution with a numerical integration solution to solve for the nearby object distortion. Second, we link the material properties and macroscopic electrical properties of objects in our solution, bridging the gap between the two, and improving the simulation capabilities of our framework.

Like [31, 32], we consider a spherical object placed in a spatially uniform, but time-varying, electric field. This is a good first approximation for including foreign objects, such as worms, in the field model because the electric field does not vary widely over the volume of the object, for small objects [32]. We aim to compute the electric field perturbation due to the object at any location in space, in particular at the locations of the mormyromast electroreceptors. To do so, we use the electric field generated by the EOD (Equation 2), measured at the location of the center of the object,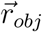. We assume that the object is small enough such that the uniform-field approximation holds. In this idealized problem, the object is placed in an spatially uniform electric field, 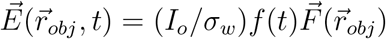. We use the dipole approximation to solve the field distortion induced by the object via Legendre series due to the azimuthal symmetry [32]. The temporal component of the EOD and the charge conservation boundary conditions make it simpler to solve the problem in the Fourier frequency domain and then invert back to the temporal domain. Let 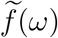 be the Fourier transform (FT) of *f*(*t*). Then, the FT of the electric potential perturbation at point 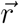 due to the spherical object of radius *a* and location 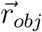 is:

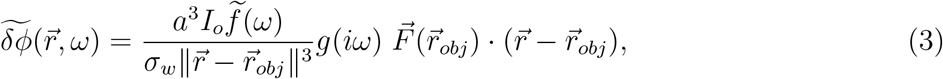

where 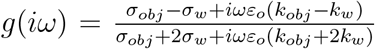, *σ*_*w*_ and *σ*_*obj*_ are the water and object conductivities, respectively, *k*_*w*_ and *k*_*obj*_ are the water and object relative permittivity constants, respectively, and *ε*_*o*_ is the vacuum permittivity. We expand *g*(*iω*) around 0, namely 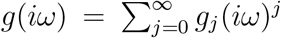. Then,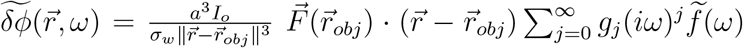. Using the inverse FT (IFT), for compactly supported functions such as *f*(*t*), we find that 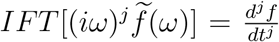. Then, the potential perturbation is:

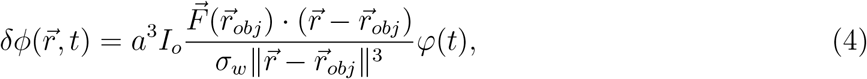

where 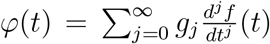. Due to the brief duration of the pulse and high frequencies of the EOD, this series does not converge for many choices of the electrical properties *σ*_*obj*_ and *k*_*obj*_. We can separate the spatial and temporal variables, thus we plug in the FT of this dipole distortion, 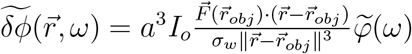 into the left hand side of Equation 3, to get:

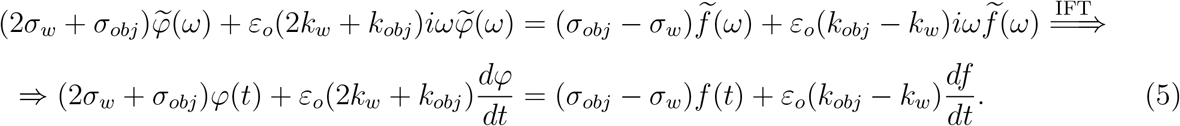

We convert from material properties to macroscopic electrical properties based on simple assumptions. We assume the spherical object with radius *a* is a resistor-capacitor object with cross section *πa*^2^ and length 2*a*, such that we estimate:

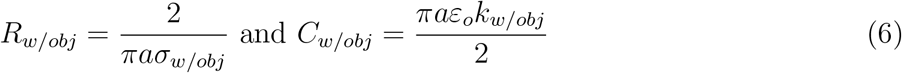

and we substitute in Equation 5 to transition between material and macroscopic object properties.

The electric field distortion is given by the spatial gradient of the potential perturbation, namely 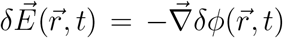. The azimuthal symmetry of the problem permits solving for the electric field perturbation using only a 2D plane which passes through the center of the object and contains 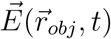. The electric field perturbation in the rest of space can be obtained by rotational symmetry. For simplicity, we translate the system such that 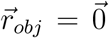. Since we already solve the temporal component, we include it here to provide the full solution:

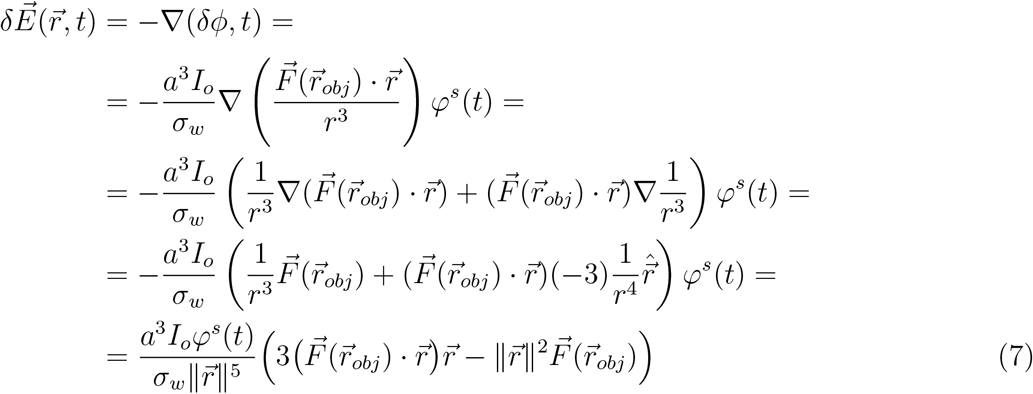

Due to the mentioned coordinate translation, 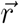 is measured from the object center. Therefore, in the simulations, we apply the 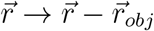 transformation.

#### A.3 Transdermal potential

The transdermal potential sensed by mormyromast receptors is given by the voltage drop across the receptor, and it is the base for the electric image formed on the skin of the fish. For a receptor at a location 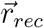 where the surface normal to the skin of the fish is given by 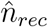, the voltage drop can be computed as:

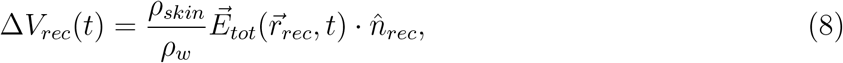

where 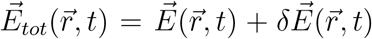 is the total electric field, due to both the EOD (Equation 2) and objects perturbations (Equation 7), if objects are present. The skin resistivity *ρ*_*skin*_ has units of Ω*m*^2^ because it is measured across the whole thickness of the skin for a surface patch, without dividing by the thickness.

